# Repurposing brewery contaminant yeast as production strains for low-alcohol beer fermentation

**DOI:** 10.1101/2021.08.01.454645

**Authors:** Kristoffer Krogerus, Ronja Eerikäinen, Heikki Aisala, Brian Gibson

## Abstract

A number of fungal isolates were recently obtained from a survey of the microbiota of multiple breweries and brewery products. Here, we sought to explore whether any of these brewery contaminants could be repurposed for beneficial use in beer fermentations, with particular focus on low-alcohol beer. 56 yeast strains were first screened for the utilization of different carbon sources, ability to ferment brewer’s wort, and formation of desirable aroma compounds. A number of strains appeared maltose-negative and produced desirable aromas without obvious off-flavours. These were selected for further scaled-up wort fermentations. The selected strains efficiently reduced wort aldehydes during fermentation, thus eliminating undesirable wort-like off-flavours, and produced a diverse volatile aroma profile. Sensory analysis of the beer samples using projective mapping identified two strains, *Trigonopsis cantarellii* and *Candida sojae*, that produced beers similar to a commercial reference lager beer. 30 L-scale wort fermentations were performed with these two strains together with a commercial *Saccharomycodes ludwigii* reference strain. Both strains performed comparably to the commercial reference, and the *T. cantarellii* strain in particular, produced low amounts of off-flavours and a significantly higher amount of the desirable monoterpene alcohol *trans*-geraniol. The strain was also sensitive to common food preservatives and antifungal compounds, and unable to grow at 37 °C, suggesting it is relatively easily controllable in the brewery, and appears to have low risk of pathogenicity. This study shows how the natural brewery microbiota can be exploited as a source of non-conventional yeasts for low-alcohol beer production.

**Take Away:**

- Fungal isolates from brewery microbiota were screened for beer production
- Numerous maltose-negative strains were tested for low-alcohol beer fermentation
- *Trigonopsis cantarellii* showed promise compared to a commercial reference strain
- *T. cantarellii* produced no off-flavours and higher levels of *trans*-geraniol

## Introduction

The global market for low-alcohol and non-alcoholic beer has increased considerably in the past decade [Statista 2021; Bellut and Arendt 2019]. This increase is driven by numerous factors, including health consciousness and growth in areas where alcohol consumption is forbidden. Compared to regular-strength beer, however, low-alcohol beer is typically afflicted with less ‘beer flavour’ and the presence of undesirable off-flavours. The flavour quality of low-alcohol beer is very much dependent on the production method, which can be broadly divided into those which are physical or biological [Brányik, Silva, Baszczyňski, Lehnert, and Almeida e Silva 2012]. The biological methods, where ethanol formation is limited, have gained much interest in the past years because of their potential for improved flavour quality. Numerous non-conventional yeasts have successfully been applied for low-alcohol beer production [Bellut, Michel, Zarnkow, Hutzler, Jacob, Atzler, Hoehnel, Lynch, and Arendt 2019; Johansson et al. 2021; Saerens and Swiegers 2013; Methner, Hutzler, Matoulková, Jacob, and Michel 2019; Capece, De Fusco, Pietrafesa, Siesto, and Romano 2021]. These yeasts lack the ability to metabolise maltose and maltotriose, the most abundant sugars in brewer’s wort, yet produce sufficient amounts of the volatile secondary metabolites characteristic of beer. Alcohol formation is therefore naturally limited to that produced from the metabolism of the wort monosaccharides. Compared to the *Saccharomyces* yeasts traditionally used in brewing, many non-conventional yeasts produce higher amounts of desirable aroma-active compounds [Gutiérrez, Boekhout, Gojkovic, and Katz 2018; Holt, Mukherjee, Lievens, Verstrepen, and Thevelein 2018; Gamero, Quintilla, Groenewald, Alkema, Boekhout, and Hazelwood 2016].

Non-conventional yeasts suitable for brewing have been isolated from a wide range of environmental niches [Cubillos, Gibson, Grijalva□Vallejos, Krogerus, and Nikulin 2019]. Repurposing of yeasts isolated from fermented food systems, like sourdoughs, kombucha and other traditional fermented beverages, has been a popular strategy [Johansson et al. 2021; Bellut et al. 2018; Tamang, Watanabe, and Holzapfel 2016]. Natural environments, like fruits, tree barks and insects, have been shown to be a good source of such strains as well [Osburn et al. 2018; Madden, Epps, Fukami, Irwin, Sheppard, Sorger, and Dunn 2018; Nikulin, Vidgren, Krogerus, Magalhães, Valkeemäki, Kangas-Heiska, and Gibson 2020; Hutzler, Michel, Kunz, Kuusisto, Magalhães, Krogerus, and Gibson 2021]. The brewery environment itself offers another potential source for non-conventional yeasts. Indeed, brewery facilities, ingredients and products are routinely tested for the presence of contaminant organisms to ensure that product quality and hygiene standards are maintained. Many yeasts, e.g. *Brettanomyces bruxellensis* and diastatic *Saccharomyces cerevisiae*, are common contaminants that can seriously influence product quality and lead to product recalls if unchecked [Tubia, Prasad, Pérez-Lorenzo, Abadín, Zumárraga, Oyanguren, Barbero, Paredes, and Arana 2018; Krogerus and Gibson 2020]. However, many of the species routinely discovered in brewery systems are not a serious threat to product quality but are rather indicative of hygiene levels [Powell and Kerruish 2017]. These benign species are commonly found in traditional fermentation systems where they are known to contribute to flavour development [Spitaels, Wieme, Janssens, Aerts, Daniel, Van Landschoot, De Vuyst, and Vandamme 2014]. Examples of such contaminants include certain *Kluyveromyces* and *Hanseniaspora* species, which are known to produce floral aromas that are considered positive in beer [Fabre, Blanc, and Goma 1998; Moreira, Mendes, Hogg, and Vasconcelos 2005], and *Torulaspora delbrueckii*, which produces fruity amyl alcohols [Canonico, Agarbati, Comitini, and Ciani 2016; Michel, Kopecká, Meier-Dörnberg, Zarnkow, Jacob, and Hutzler 2016].

A recent survey of the microbiota in multiple breweries yielded a large collection of fungal isolates [Sohlberg, Sarlin, and Juvonen 2021]. While a broad microbiota may typically be seen as a negative in an industrial brewery, we wanted to turn this into a positive. The aim was to first use high-throughput screening to identify maltose-negative strains suitable for low-alcohol brewing. A subset of isolates was further screened in small-scale and 2 L-scale wort fermentations, after which chemical and qualitative sensory analysis was performed on the resulting beers. Following this screening, two candidate strains were selected for scaled-up wort fermentations and descriptive sensory analysis. The present study highlights how the natural brewery microbiota can be exploited for low-alcohol beer production.

## Materials & Methods

### Yeast strains

A list of all non-conventional yeast strains included in this study is found in Supplementary Table S1. In addition, the following three strains from VTT’s culture collection (http://culturecollection.vtt.fi) were included as commercial references: *Saccharomyces cerevisiae* VTT-A75060, *Saccharomyces pastorianus* VTT-A63015, and *Saccharomycodes ludwigii* VTT-C181010.

### Microplate cultivations

Growth on various carbon sources and in the presence of 100 mg/L hop-derived iso-alpha acids was tested in microplate cultivations. The microcultures were carried out in 100-well honeycomb microtiter plates at 25 °C (with continuous shaking), and their growth dynamics were monitored with a Bioscreen C MBR incubator and plate reader (Oy Growth Curves Ab, Finland). The wells of the microtiter plates were filled with 300 µL of defined medium (0.67% yeast nitrogen base without amino acids, 1% carbon source). Precultures of the strains were started in 20 mL YPD medium (1% yeast extract, 2% peptone, 2% glucose) and incubated at 25 °C with shaking at 120 rpm overnight. The optical density at 600 nm was measured, and precultures were washed and resuspended to an OD600 value of 3. The microcultures were started by inoculating the microtiter plates with 10 µL of cell suspension per well (for an initial OD600 value of 0.1) and placing the plates in the Bioscreen C MBR. The optical density of the microcultures at 600 nm was automatically read every 30 min. Four replicates were performed for each strain in each medium. Growth curves for the microcultures were modelled based on the OD600 values over time using the ‘GrowthCurver’-package for R.

The ability to produce phenolic off-flavour was estimated using the absorbance-based method described by Mertens et al. (2017).

### Wort fermentations

Small-scale fermentations were carried out in 100 mL Schott bottles capped with glycerol-filled airlocks. Yeast strains were grown overnight in 25 mL YPD medium at 25 °C. The pre-cultured yeast was then inoculated into 80 mL of all-malt wort (extract ranged from 5 to 10 °Plato) at a rate of 1 to 2.5 g fresh yeast L^−1^ (depending on °Plato of wort). Fermentations were carried out in duplicate at 25 °C for 7 days. Fermentations were monitored by mass lost as CO_2_.

2 L-scale fermentations were carried out in 3 L cylindroconical stainless steel fermenting vessels containing 2 L of 5 °P wort. Yeast was propagated in autoclaved wort. The 5 °P wort (23.3 g maltose, 6.3 g maltotriose, 5.8 g glucose, and 1.6 g fructose per litre) was produced at the VTT Pilot Brewery from barley malt. The wort was oxygenated to 10 mg L^−1^ prior to pitching (Oxygen Indicator Model 26073 and Sensor 21158; Orbisphere Laboratories, Switzerland). Yeast was inoculated at a rate of 1 g fresh yeast L^−1^. The fermentations were carried out in duplicate at 25 °C for 7 days.

30L-scale fermentations were carried out in 40 L stainless steel tanks containing 30 L of 7.5 °P wort. The 7.5 °P wort (36.3 g maltose, 10.1 g maltotriose, 9.7 g glucose, and 2.4 g fructose per litre) was produced at the VTT Pilot Brewery from barley malt. Yeast was propagated and inoculated as above. Fermentations were carried out for 7 days, after which the beer was collected in sterilized kegs for maturation. Green beers were matured at 10 °C for five days before five days’ stabilization at 0 °C, after which time the beers were depth filtered (Seitz EK filter sheets with a relative retention of <1.0 μm, Pall Corporation, USA), carbonated to 5 g/l of CO2 and bottled. The bottled beer was further pasteurized at 60 °C for 30 minutes.

### Chemical analysis

The alcohol content of the final beer was measured with an Anton Paar density meter DMA 5000 M with Alcolyzer beer ME and pH ME modules (Anton Paar GmbH, Austria).

Volatile aroma compounds were analysed using headspace solid phase micro-extraction coupled with gas chromatography (Agilent 7890A) - mass spectrometry (Agilent 5975C; HS-SPME-GC-MS) by modifying the method used by Rodriguez-Bencomo et al. [Rodriguez-Bencomo, Muñoz-González, Martín-Álvarez, Lázaro, Mancebo, Castañé, and Pozo-Bayón 2012]. 6 mL of beer sample, 1.8 g of NaCl, and 50 µL of internal standard solution (containing 1.28 µg 3-octanol, 1.19 µg 3,4-dimethylphenol) were added to 20 mL headspace vials. The samples were pre-incubated in Gerstel MPS autosampler at 44.8 □ for 10 min and the volatiles were extracted by using a 2 cm divinylbenzene/ carboxen/polydimethylsiloxane (DVB/CAR/PDMS) fibre (Supelco) at 44.8 □ for 46.8 min. The samples were injected in splitless mode (10 min desorption time at 270 □) and the compounds were separated on an HP-Innowax silica capillary column (60 m, 0.250 mm i.d., 0.25 µm film thickness). The oven temperature program was from 40□°C (3□min) to 240□°C (4□°C□min^−1^) and the final temperature was held for 15□min. The MS data were collected at a mass range of 35-450 amu. Identification was based on spectral data of reference compounds and those of NIST 08 library. Calibration curves determined for 2-methoxy-4-vinylphenol, 2-phenylethyl acetate, α-terpineol, β-citronellol, *cis*-geraniol, *trans*-geraniol, isopentyl acetate, linalool, and ethyl esters of acetic, butyric, decanoic, hexanoic and octanoic acids, respectively (r^2^= 0.933-0.999). Other compounds were quantified by using internal standards (3-octanol or 3,4-dimethylphenol).

Aldehydes were analysed as oximes by using a headspace sampler (Agilent 7697A) coupled with gas chromatograph (Agilent 7890B) and compounds were detected using a Micro Electron Capture Detector (HS-GC-ECD). Carbonyl compound standards were 2-methylpropanal, 2-methylbutanal, 3-methylbutanal, hexanal, furfural, methional, phenylacetaldehyde, and (*E*)-2-nonenal (Aldrich, Finland). A stock solution containing a mixture of the standard compounds in ethanol was prepared at 1000□μg□L^−1^ each. The calibration range was 0.5–40□μg□L^−1^ and dilutions were prepared in 5% ethanol. The sum of the peak areas of the two geometrical isomers (*E* and *Z*) was used for calculations. An aqueous solution of derivatization agent O-(2,3,4,5,6-pentafluorobenzyl)-hydroxylamine (PFBOA) (Sigma-Aldrich) was prepared at a concentration of 6□g/L. One hundred microliters of this solution and 5□mL of deionized water or beer were placed in a 20 mL glass vial and sealed with a crimp cap (Agilent). The sample/standard vial was then placed in the headspace sampler with following conditions: sample equilibrium in oven for 30□min□at 60□°C, after which 1□min of injection of sample fill pressurized at 25 psi. Loop temperature was 100□°C and transfer line was held at 110□°C. The following GC conditions were applied: HP-5 capillary column, 50□m□×□0.32□mm ×1.05□μm (J&W Scientific, Folsom, CA). Helium was the carrier gas at a flow rate of 1.0□mL/min and for ECD, nitrogen make up gas was applied at a flow rate of 30□mL/min. The front inlet temperature was 250□°C. The injection was in the split mode and the split ratio 10:1 was applied. The oven temperature program used was 40□°C for 2□min, followed by an increase of 10□°C□min^−1^ to 140□°C (held 5□min) and 7□°C□min^−1^ to 250□°C. The final temperature was held for 3□min.

Diacetyl, dimethyl sulphide, higher alcohol and ester concentrations in the beers produced at 30L-scale were analysed at Campden BRI (UK).

### Sensory analysis

Projective Mapping with Ultra Flash profiling was conducted for the beers based on established methods [Risvik, McEwan, Colwill, Rogers, and Lyon 1994; Perrin and Pagès 2009; Aisala, Laaksonen, Manninen, Raittola, Hopia, and Sandell 2018] and carried out in an ISO-8589 sensory evaluation laboratory at VTT. A total of 10 selected and trained assessors participated in the evaluation. Written informed consents were obtained from the participants prior to the evaluation. A commercial beer sample was included in the samples twice (second time as a blind duplicate) for a total number of 9 samples. The samples were served in black beer glasses covered in lids, marked with 3-digit codes and served in random order. The evaluation was done in two parts: the first Projective mapping based on odour properties and the second one based on taste and flavour properties. An A3 size paper was used for placing the samples. The assessors were asked to first smell or taste the samples and make notes on the provided A4 paper with instructions. Then they were asked to physically place the samples on the A3 evaluation paper so that the similarity of the samples was reflected in their distances. After the assessor was ready with the placement, they were asked to mark the locations of the samples with the 3-digit code, mark possible groups, and write descriptive terms next to the samples or sample groups. The filled papers were scanned, and the coordinates of each sample were measured digitally using the ImageJ software [Schneider, Rasband, and Eliceiri 2012]. The descriptive terms were collected into a contingency table. The resulting dataset was analysed with Multiple Factor Analysis (MFA) using R and the ‘SensoMineR’ package [Le and Husson 2008].

The descriptive profiling of the three beer samples was conducted by nine trained assessors in the same sensory evaluation laboratory as above. The base attribute list was developed by four trained assessors in a consensus session. After that, the whole panel was trained in two groups where they refined this base sensory lexicon, discussed the intensity ranges of the samples, and decided on reference samples for the intensities. The final sensory profile had 6 odour attributes, 4 taste and flavour attributes and 2 mouthfeel attributes. The samples (50 mL) were served in black beer glasses with 3-digit codes and the serving order was randomized with a Latin squares design. The attribute intensities were evaluated with a 0–10 continuous line scale that was anchored with 0 = attribute non-perceivable, and 10 = attribute perceived as very intense. The data were collected with Compusense five 5.6 (Compusense Inc., Guelph, ON, Canada). Water was used as the palate cleanser and the assessors were instructed to spit the sample out after tasting. The descriptive sensory data was analysed with a two-way mixed model analysis of variance with samples as the fixed factor and the assessors as a random factor using SPSS version 26 (IBM Corp, Armonk, NY, USA). Tukey’s HSD was used as the post hoc test.

### Assessment of process and health safety

The tolerance to common food preservatives was tested in microplate cultivations using the BioScreen C incubator similarly as described above. Cultivations were carried out in 150 µl of YPD (1% glucose w/v) supplemented with 5% ethanol, 150 mg/L sodium benzoate (Sigma-Aldrich, Darmstadt, Germany), 250 mg/L potassium sorbate (Sigma-Aldrich, Darmstadt, Germany), or 200 mg/L potassium metabisulfite (Brown, Poland). All media were adjusted to pH 4 with HCl.

Biofilm formation was assessed by crystal violet staining of microplate cultures in 96-well plates [Shukla and Rao 2017]. Wells were filled with 250 µL of 10 °Plato wort, and inoculated with 2.5 µL of overnight culture in five replicates. Plates were statically incubated for five days at 20 °C. The yeast cultures were removed by pipetting, and the plate was rinsed with sterile deionized water. 300 µL of 0.1 % crystal violet solution was placed in the wells for five minutes, after which the plate was rinsed three times with sterile deionized water. The plate was left to air-dry for 15 minutes in a laminar flow cabinet. The remaining crystal violet, which was still bound to the cells, was dissolved with 300 µl of 95 % ethanol. Absorbances of the wells were measured at 595 nm with a Multiskan EX plate reader (Labsystems Oy, Finland). The plate was left overnight at 4 °C and absorbance was re-measured.

Tolerance to nine antifungal compounds was tested using Sensititre YeastOne YO10 plates (Thermo Scientific, Finland) using kit instructions. Concentrations of biogenic amines in beers was measured at Campden BRI (UK).

### Whole-genome sequencing

The whole-genome of *Trigonoposis cantarellii* P-69 was sequenced at NovoGene (UK). DNA was extracted using the method described by Denis et al. [2018]. Sequencing was carried out on a NovaSeq 6000 instrument (Illumina). The 150 bp paired-end reads have been submitted to NCBI-SRA under BioProject number PRJNA748016. Reads were trimmed and filtered with fastp using default settings (version 0.20.1; [Chen, Zhou, Chen, and Gu 2018]). Trimmed reads were assembled with SPAdes (version 3.10.1; [Bankevich et al. 2012]). The resulting assembly was filtered using CVLFilter (using minimum length of 500 and minimum coverage of 10) to remove low-coverage contigs resulting from potential contaminating DNA [Douglass, O’Brien, Offei, Coughlan, Ortiz-Merino, Butler, Byrne, and Wolfe 2019]. BUSCO (Benchmarking Universal Single-Copy Orthologs; version 3.0.2; using the saccharomycetales_odb9 dataset) was used to assess the gene set completeness of the assembly [Waterhouse, Seppey, Simão, Manni, Ioannidis, Klioutchnikov, Kriventseva, and Zdobnov 2018].

The *T. cantarellii* P-69 assembly was compared to the assemblies of other closely related species obtained from NCBI-Assembly under BioProject PRJNA429441 [Shen et al. 2018]. A multiple sequence alignment of BUSCO genes was performed as described in [Steenwyk, Shen, Lind, Goldman, and Rokas 2019]. In brief, the amino acid sequence of orthologous single-copy BUSCO genes with >50% taxon occupancy were extracted (using genomic_BUSCOs2uscofa.py from https://github.com/JLSteenwyk/Phylogenetic_scripts) and aligned with MAFFT (version 7.407; [Katoh and Standley 2013]). The alignments were trimmed with trimal (version 1.4.rev15; [Capella-Gutierrez, Silla-Martinez, and Gabaldon 2009]), and a maximum likelihood phylogenetic tree was constructed using IQ-TREE (version 1.5.5; [Nguyen, Schmidt, Von Haeseler, and Minh 2015]) with the JTT+I+G4 model and 1000 bootstrap replicates [Minh, Nguyen, and Von Haeseler 2013]. In addition, a neighbour-joining tree based on MinHash distances [Ondov, Treangen, Melsted, Mallonee, Bergman, Koren, and Phillippy 2016] was generated using mashtree [Katz, Griswold, Morrison, Caravas, Zhang, Bakker, Deng, and Carleton 2019]

## Results and Discussion

### Phenotypic screening of fungal brewery isolates for brewing-relevant traits

A total of 56 yeast strains, previously isolated from industrial brewery environments [Sohlberg, Sarlin, and Juvonen 2021], were screened for various brewing-relevant phenotypes. Among the strains were 18 different species from 10 genera, and the isolation environments included brewery air, brewery surfaces, raw materials, and spoiled products. A *Saccharomyces cerevisiae* ale strain, *Saccharomyces pastorianus* lager strain, and *Saccharomycodes ludwigii* low-alcohol strain were included as references. First, their growth on various carbon sources (glucose, fructose, maltose, and maltotriose) was measured in microplate cultivations (Figure 1A). The strains could be broadly grouped into three groups: those capable of growing on both maltose and maltotriose, those only on maltose, and those on neither. For full-strength beer fermentation, a yeast strain must be capable of utilizing at least maltose (constituting approx. 60% of the fermentable wort sugars), and preferably maltotriose as well (constituting approx. 20% of the fermentable wort sugars). Maltose-negative strains, on the other hand, have shown promise for low-alcohol beer fermentations [Johansson et al. 2021; Bellut et al. 2019]. In addition, growth in the presence of 100 mg/L hop-derived iso-alpha acids was tested, and no inhibition was observed for any of the strains (data not shown).

**Figure 1.**
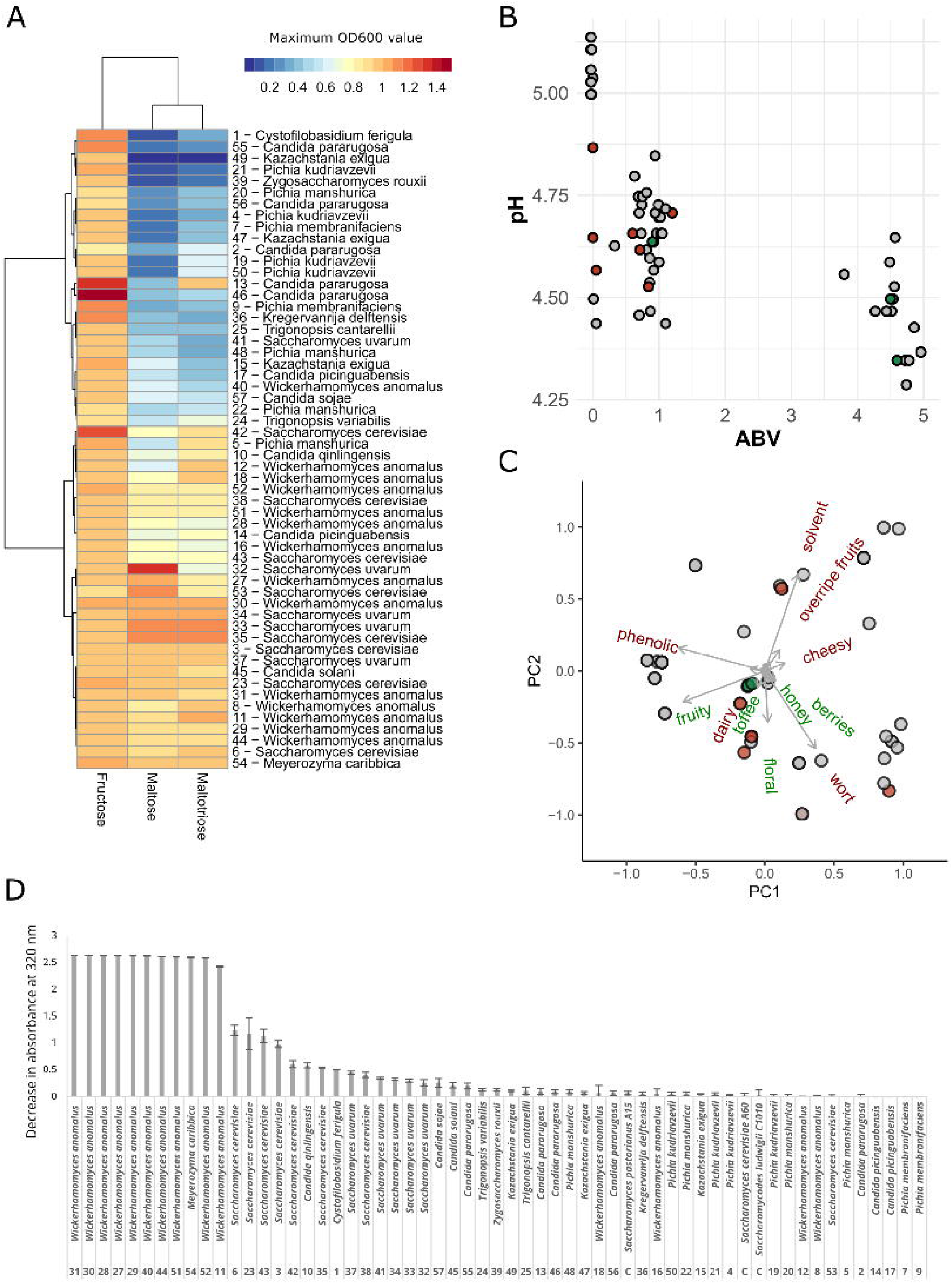
Phenotypic screening of 56 strains isolated from brewery environments. (**A**) The ability to grow on fructose, maltose and maltotriose as sole carbon sources. (**B**) The relationship between beer pH and alcohol by volume after wort fermentation with the 56 strains and three controls. The controls are marked with green, while the strains selected for further characterization are marked with red. (**C**) Principal component analysis of the results from the sniffing of beers produced with the 56 strains and three controls. Undesirable aromas are coloured red, while desirable aromas green. Points are coloured as in (**B**). (**D**) The ability to decarboxylate ferulic acid (to 4-vinyl guaiacol) among the 56 strains and three controls as assessed by the decrease in absorbance at 320 nm after growth in media supplemented with 100 mg/L ferulic acid.

Small-scale anaerobic wort fermentations were also carried out with all the strains to confirm the results observed during the microplate cultivations, and to test which strains produce pleasant aromas. Again, the strains grouped into three main groups: those producing approx. >4% ABV, approx. 1% ABV, and those showing negligible alcohol formation (Figure 1B). Expectedly, *Saccharomyces* yeasts performed well in wort and grouped into the first group. The third group consisted of multiple strains showing no signs of fermentation in wort. Interestingly, however, a pH decrease was observed for five strains, indicating metabolic activity despite no ethanol formation.

The fermented worts were also subjected to a ‘sniff’ test, to identify strains producing pleasant or clearly undesirable aromas (Figure 1C). Undesirable solvent-like aromas were detected in a large number of strains, particularly among *Wickerhamomyces anomalus* isolates. Undesirable wort-like aromas were also detected for many strains, mainly those performing poorly during wort fermentations. Desirable fruity aromas were observed mainly among the *Saccharomyces* strains, however, they were frequently coupled with undesirable phenolic aromas. The ability to decarboxylate ferulic acid to 4-vinylguaiacol (i.e. phenolic off-flavour formation) was therefore tested for all strains (Figure 1D), and found in approx. 45%. Out of the 56 strains that were screened, nine had a pleasant aroma lacking any obvious off-flavours (e.g. phenolic, solvent-like, or wort-like). These included strains of *Candida pararugosa, Candida sojae, Kregervanrija delftensis, Pichia manshurica, S. cerevisiae, Trigonopsis cantarellii, Trigonopsis variabilis*, and *Zygosaccharomyces rouxii*. As the majority of these were maltose-negative, we next explored whether they could be used for low-alcohol beer fermentation.

### Identifying strains suitable for low-alcohol beer fermentations

Seven brewery isolates (Table 1), as well as a commercial *S. ludwigii* reference strain, were selected for further trials following the initial pre-screening. 80 mL-scale wort fermentations were first performed. With the exception of those fermented with the *W. anomalus* isolate P-2.4, the alcohol content of the beers remained below 0.70% alcohol by volume (Table 1). pH values of the beers ranged from 4.4 to 5.1 (dropping from 5.3 in the wort), with the highest values measured in the beer fermented with the *S. ludwigii* reference strain. A beer pH value below 5 is typically recommended for microbiological safety reasons [Menz, Aldred, and Vriesekoop 2011]. The main contributors to the wort-like flavour of unfermented wort, a flavour that is considered undesirable in beer, are a number of unbranched and branched-chain aldehydes. Hence, we measured the concentration of six key aldehydes in the beers. The wort aldehydes were efficiently reduced by all strains (Figure 2A), and were in many cases below the levels measured in a commercial full-strength lager beer (dashed line). Methional, in particular, with its low flavour threshold of 5 µg/L, is considered one of the main causes of wort-like flavour [Gernat, Brouwer, and Ottens 2020]. Here, the majority of the strains reached concentrations below the flavour threshold. Volatile aroma compounds in the beers were also measured with HS-SPME-GC/MS. The aroma profiles could be broadly divided into four groups (Figure 2B), with the wort and the beer fermented with *W. anomalus* P-2.4 forming two outliers. The wort sample was expectedly characterized by aldehydes, while the *W. anomalus* beer was characterized by the highest concentrations of volatile esters (Figure 2C). *Meyerozyma caribbica* T-31 and *Candida sojae* T-39 clustered together with the *S. ludwigii* reference strain, while the other strains clustered intermediate between the three other groups.

**Table 1.** Alcohol by volume (%) and pH of beers produced with 7 brewery isolates and a *S. ludwigii* reference at lab-scale. Standard deviation in parenthesis.

| Strain | Alcohol by volume (%) | pH |
| --- | --- | --- |
| *Wickerhamomyces anomalus* P-2.4 | 2.13 ($\pm 0.03$) | 4.57 ($\pm 0.08$) |
| *Pichia manshurica* P-5.2 | 0.04 ($\pm 0.01$) | 4.61 ($\pm 0.03$) |
| *Trigonopsis variabilis* P-5.8 | 0.02 ($\pm 0.01$) | 4.78 ($\pm 0.01$) |
| *Trigonopsis cantarellii* P-69 | 0.58 ($\pm 0.04$) | 4.40 ($\pm 0.03$) |
| *Kregervanrija delftensis* R-7 | 0.10 ($\pm 0.01$) | 4.36 ($\pm 0.01$) |
| *Meyerozyma caribbica* T-31 | 0.61 ($\pm 0.01$) | 4.52 ($\pm 0.04$) |
| *Candida sojae* T-39 | 0.50 ($\pm 0.00$) | 4.66 ($\pm 0.01$) |
| *Saccharomycodes ludwigii* C181010 | 0.70 ($\pm 0.00$) | 5.09 ($\pm 0.02$) |

**Figure 2.**
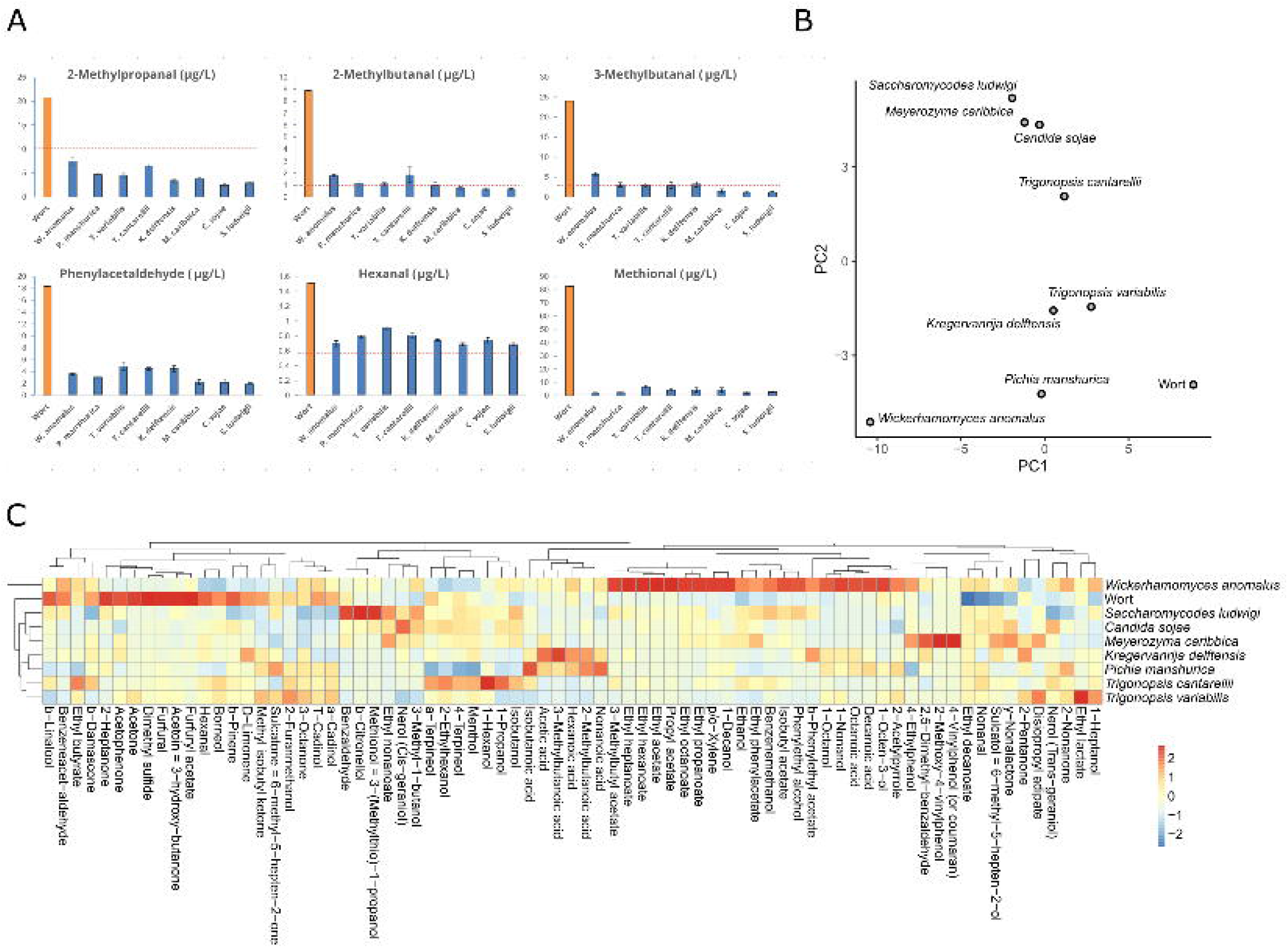
(**A**) Concentrations of aldehydes in wort and beers (µg/L). The dotted line indicates concentrations in a commercial pale lager beer. (**B**) Principal component analysis (PC1 vs PC2) of the concentrations of volatile aroma compounds detected in the wort and beers. (**C**) Heatmap of the concentrations of volatile aroma compounds detected in the wort and beers. The heatmap is coloured based on Z-scores of the concentrations for each compound (blue: negative Z-score, red: positive Z-scores).

Based on these results, five brewery isolates (*C. sojae, K. delftensis, T. cantarellii, T. variabilis*, and *W. anomalus*) and the *S. ludwigii* reference strain were selected for 2 L-scale wort fermentations and subsequent sensory analysis. The alcohol content of the beers ranged from 0.0% to 0.6% alcohol by volume after one week of fermentation, while beer pH again ranged from 4.5 to 5.1 (Table 2). The beer was collected for sensory analysis, which was carried out by ten trained participants at VTT’s sensory lab. We wanted to identify what beers were most similar to a commercial full-strength lager beer, and for that sensory analysis was performed using Projective Mapping with Ultra Flash Profiling [Risvik, McEwan, Colwill, Rogers, and Lyon 1994; Perrin and Pagès 2009]. Here, the six experimental beer samples, along with two duplicate commercial lager beers and a wort sample, were assessed based on odour and flavour. The results were analysed with multiple factor analysis (MFA), and they revealed that the experimental beers were mostly similar, and tended to group between the wort sample and the commercial beer (Figure 3A). The *W. anomalus* beer was again a clear outlier, as the odour and flavour were dominated by solvent-like tones. Based on flavour, the beers fermented with *T. cantarellii* and the *S. ludwigii* reference strain were scored closest to the commercial full-strength lager (Figure 3B). Some common odour and flavour descriptors for these two strains and *C. sojae* were ‘mild’, ‘fruity’, and ‘sweet’ (data not shown). On the other hand, *T. variabilis, K. delftensis* as well as *S. ludwigii* were described as ‘worty’. Based on the fermentation and sensory results, *T. cantarellii* and *C. sojae* were chosen for pilot-scale fermentations.

**Table 2.** Alcohol by volume (%) and pH of beers produced with 5 brewery isolates and a *S. ludwigii* reference at 2L-scale. Standard deviation in parenthesis.

| Strain | Alcohol by volume (%) | pH |
| --- | --- | --- |
| *Wickerhamomyces anomalus* P-2.4 | 0.60 ($\pm 0.00$) | 4.64 ($\pm 0.01$) |
| *Trigonopsis variabilis* P-5.8 | 0.02 ($\pm 0.02$) | 4.54 ($\pm 0.42$) |
| *Trigonopsis cantarellii* P-69 | 0.14 ($\pm 0.00$) | 4.81 ($\pm 0.01$) |
| *Kregervanrija delftensis* R-7 | 0.00 ($\pm 0.00$) | 5.06 ($\pm 0.08$) |
| *Candida sojae* T-39 | 0.22 ($\pm 0.01$) | 4.72 ($\pm 0.03$) |
| *Saccharomycodes ludwigii* C181010 | 0.43 ($\pm 0.00$) | 4.84 ($\pm 0.04$) |

**Figure 3.**
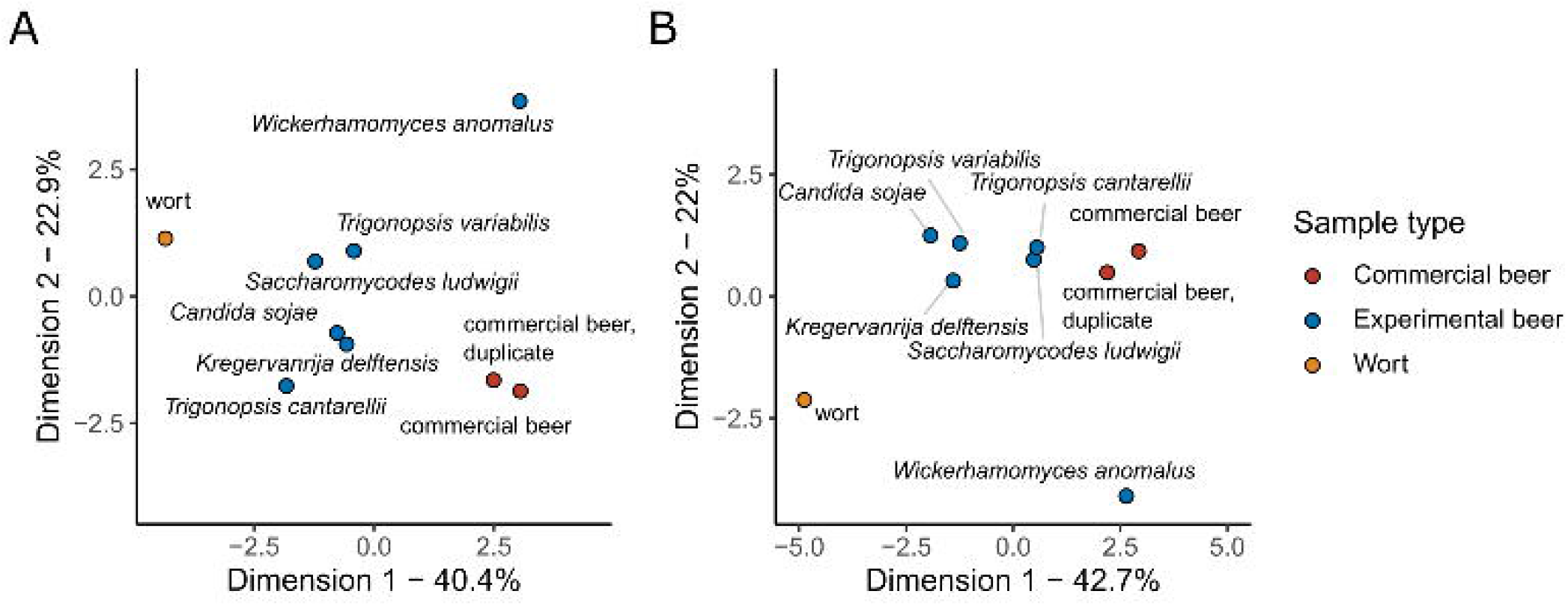
Sensory analysis of beers made with selected isolates. Sensory analysis was performed using projective mapping based on (**A**) odour and (**B**) flavour. Experimental beers were compared to an unfermented wort sample and two replicates of a commercial full-strength lager beer.

### Pilot-scale low-alcohol wort fermentations with selected strains

30 L-scale wort fermentations were carried out with brewery isolates *T. cantarellii* and *C. sojae. S. ludwigii* was again included as a reference strain. Fermentations progressed similarly to previous smaller-scale fermentations, with the beers reaching alcohol levels of 0.29%, 0.48%, and 0.68% alcohol by volume, respectively, after one week (Figure 4A). pH values in all three beers were identical (4.8). Fermentation progressed the fastest by *S. ludwigii*. Following fermentation, samples were drawn for chemical analysis, after which beers were carbonated and bottled for sensory analysis.

**Figure 4.**
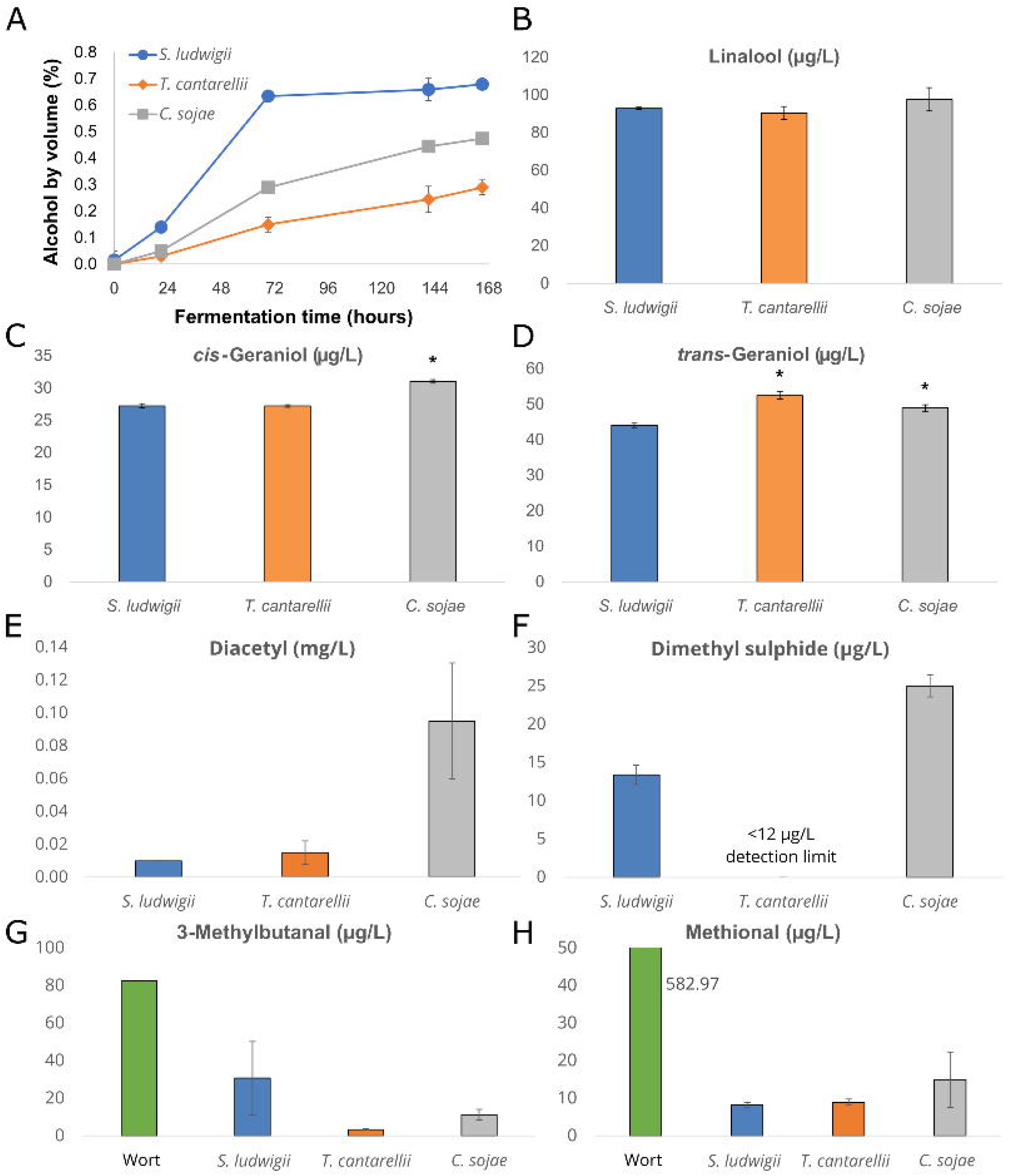
Fermentation profiles and chemical analysis of beers produced at 30L-scale. (**A**) Alcohol by volume (%) in the fermenting wort. Concentrations of (**B**) linalool, (**C**) *cis-*geraniol, (**D**) *trans*-geraniol, (**E**) diacetyl, (**F**) dimethyl sulphide, (**G**) 3-methylbutanal, and (**H**) methional in the beers and wort. All values are the mean of two biological replicates. Error bars when present show standard deviation. (**C**) and (**D**) An asterisk indicates a significant difference (*p<*0.05) compared to the *S. ludwigii* reference strain as determined by an unpaired two-tailed t-test.

A number of yeast-derived aroma-active esters play a central role in the aroma of beer [Pires, Teixeira, Brányik, and Vicente 2014], however, the concentration of these is typically low in beer fermented with maltose-negative yeast from reduced metabolism [Bellut and Arendt 2019]. Here, the beers also showed low amounts of esters and higher alcohols, with values in many cases being below detection limits (Table 3). The highest concentrations were measured in the beer produced with the *S. ludwigii* reference strain. HS-SPME-GC/MS analysis was used to measure other volatile aroma compounds as well. Differences in monoterpene alcohol concentrations were observed in the beers. Levels of the hop-derived linalool were equal in all beers, however, *trans-*geraniol and *cis-* geraniol levels were significantly higher in the *C. sojae* beer, while *trans-*geraniol levels were significantly higher in the *T. cantarellii* beer (Figure 4B-D). These monoterpenes contribute floral and citrus-like aroma to the beer. *cis-*geraniol levels were below reported flavour thresholds (80 µg/L; [Takoi, Itoga, Koie, Kosugi, Shimase, Katayama, Nakayama, and Watari 2010]), but sub-threshold concentrations may nevertheless contribute positively to the beer aroma through synergistic effects with other monoterpenes [Dietz, Cook, Huismann, Wilson, and Ford 2020]. The varying concentrations of monoterpene alcohols in the beers suggests potential biotransformation of hop-derived monoterpene alcohols [Serra Colomer, Funch, Solodovnikova, Hobley, and Förster 2020].

**Table 3.**
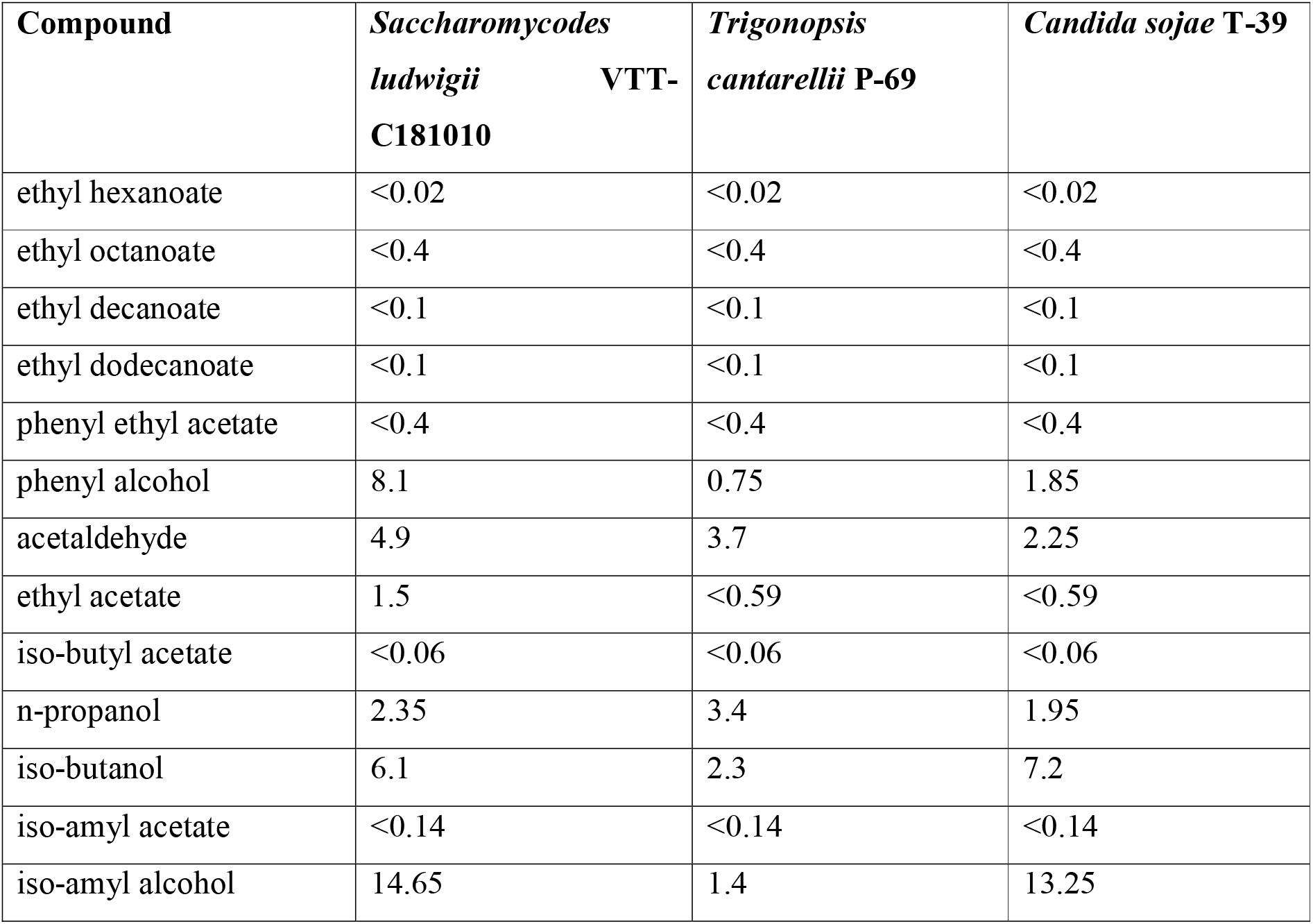
Concentrations of higher alcohols and esters (mg/L) in the beers produced at 30L-scale.

In regards to off-flavours, aldehyde levels in all beers again decreased from those measured in the wort (Figure 4G-H). Lowest aldehyde levels were measured in the beer fermented with *T. cantarellii*. Beers fermented with *S. ludwigii* tended to have higher levels of branched-chain aldehydes. Methional concentrations again decreased substantially in the beers compared to the levels in the wort. In addition to the aldehydes, concentrations of both diacetyl and dimethyl sulphide close to their respective flavour thresholds were observed in the *C. sojae* beers (Figure 4E-F). Diacetyl, with a butter-like aroma, is a common off-flavour in beer [Krogerus and Gibson 2013]. It is both indirectly generated by and reduced by the yeast during fermentation, and high beer concentrations are typically indicative of a too short fermentation time. Dimethyl sulphide, with a ‘cooked corn’ aroma, is considered undesirable at concentrations above 100 µg/L [Anness and Bamforth 1982]. Concentrations of these two off-flavours were considerably below the flavour threshold in the other two beers.

The bottled beers were subjected to sensory analysis at VTT’s sensory lab. Nine trained participants performed descriptive profiling of 12 attributes, and a commercial non-alcoholic lager beer was included as a reference. The three experimental and one reference beer scored similarly across the 12 attributes (Figure 5). Statistically significant differences were observed only for sweetness, which divided the samples into two groups: the more intense sweetness of *T. cantarellii* and *C. sojae* beers, while the reference beer and *S. ludwigii* beers were less sweet. Of the experimental beers, the one fermented with the commercial *S. ludwigii* strain scored lowest in fruitiness, possibly from the lower concentrations of monoterpene alcohols. As already revealed during chemical analysis, diacetyl notes were also detected by the panel in the *C. sojae* beer. The other beers were free of obvious off-flavours. Based on the pilot-scale fermentations, along with subsequent sensory and chemical analysis, *T. cantarellii* appears to be a promising candidate for low-alcohol beer fermentation.

**Figure 5.**
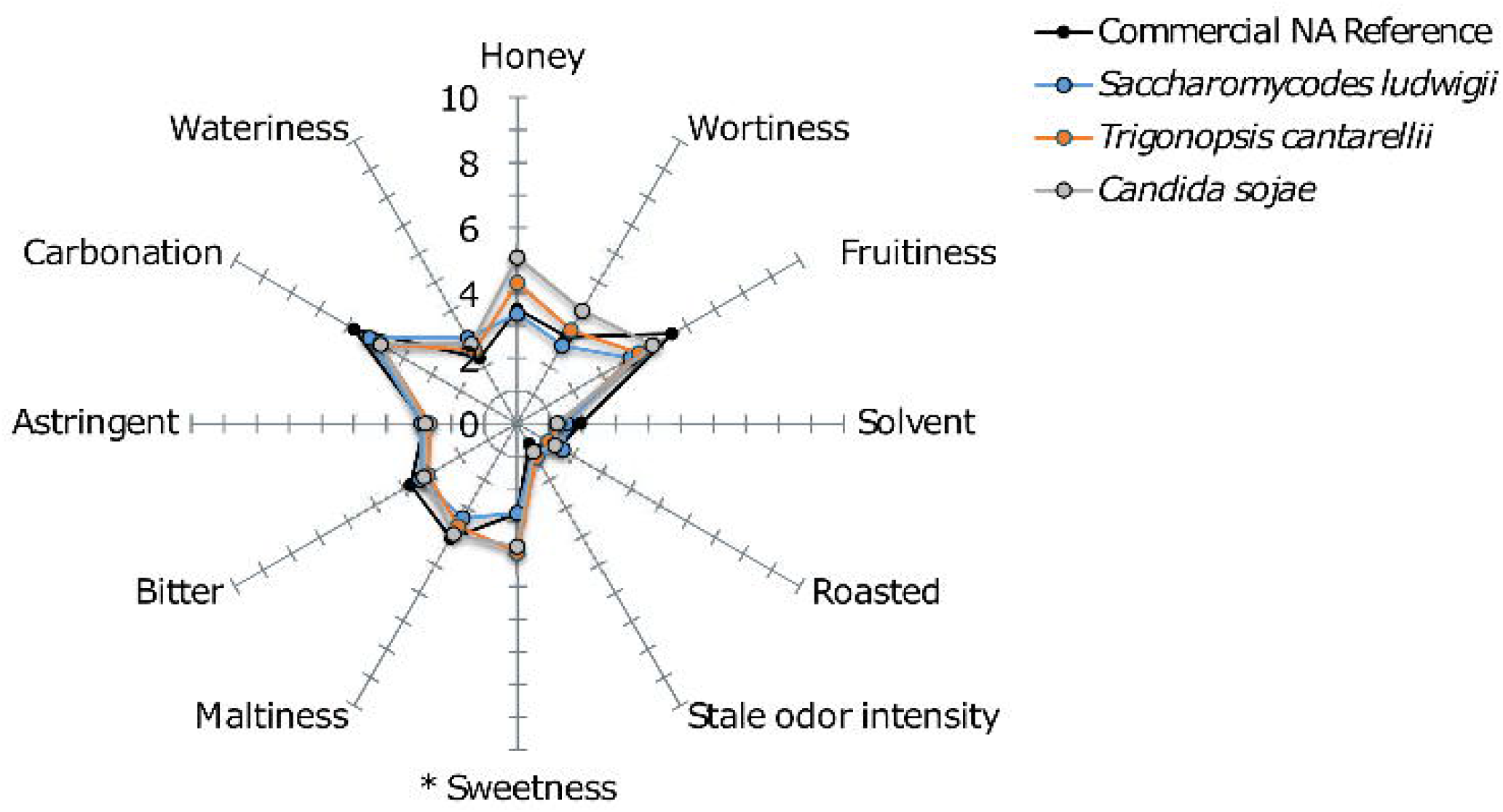
Descriptive flavour profiling of beers produced at 30L-scale. Sweetness is the only attribute with statistically significant difference (*p* < 0.05, marked with an asterisk) between samples in the two-way mixed model ANOVA.

### Process and hygiene control

As many non-conventional yeasts have not been used in food or beverage production before, it is vital that they are safe to use from both a health and process hygiene perspective. Particularly, since the yeasts studied here were already isolated from the brewery environment. To assess process safety, the tolerance of *T. cantarellii* and *C. sojae* to common food preservatives [Kregiel 2015], as well as potential for biofilm formation, was tested (Figure 6). Modern breweries often also produce non-beer beverages in the same facilities, which increases the need for controlling potential cross-contaminations. *T. cantarellii* was inhibited by all of the tested preservatives, while *C. sojae* was able to grow in the presence of 150 mg/L benzoate and 200 mg/L sulphite (Figure 6A and B). Biofilm potential was estimated using crystal violet staining of washed microculture plates, and *T. cantarellii* produced significantly less biofilm than the *S. pastorianus* A-63015 control (Figure 6C). *C. sojae*, on the other hand, produced significantly more biofilm than the control. Hence, *T. cantarellii* appears to be relatively easy to control from a process hygiene perspective, as it can be controlled using typical food preservatives in non-beer beverages, and shows little biofilm potential.

**Figure 6.**
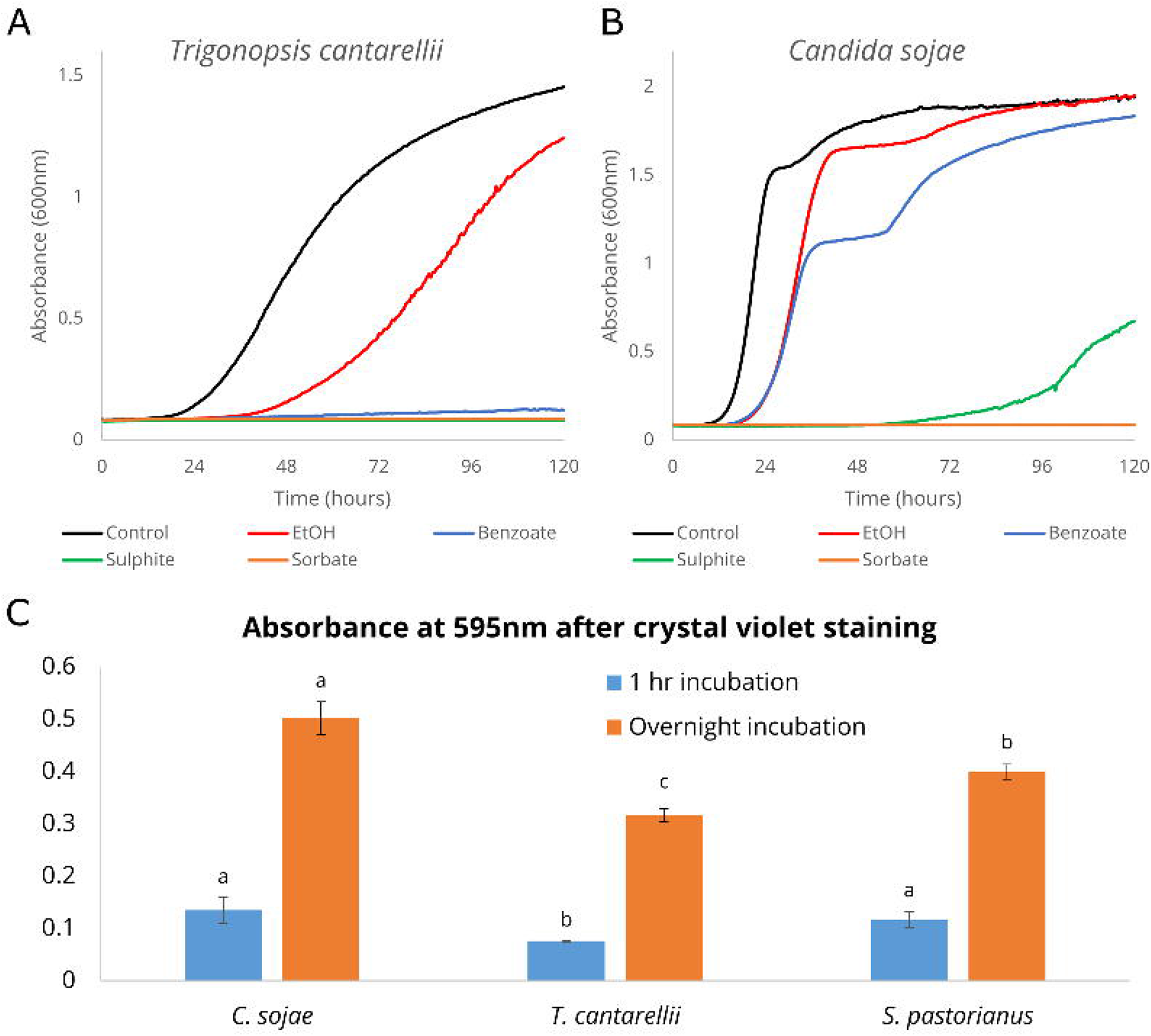
Growth of (**A**) *T. cantarellii* P-69 and (**B)** *C. sojae* T-39 in the presence of common food preservatives. (**C**) Biofilm formation after cultivation in wort for 5 days. Different letters in the two series indicate significant differences (*p<*0.05) as determined by one-way ANOVA and Tukey’s posthoc test.

To assess the potential health risks associated with the *T. cantarellii* and *C. sojae* isolates, a series of tests were carried out. First, the tolerance of the *T. cantarellii* and *C. sojae* isolates to nine antifungals was tested, to ensure that any emerging infection would be treatable. *C. sojae* was sensitive to all of the tested antifungals, while *T. cantarellii* was only able to grow in the presence of 5-fluorocytosine at the tested concentrations (MIC values in Table 4). Neither of the strains were able to grow at 37 °C (i.e. body temperature), suggesting a low potential for pathogenicity. Furthermore, the strains only produced trace levels of biogenic amines, and concentrations remained below those found in commercial beer [Poveda 2019]. Both isolates therefore appear to be relatively safe from a health perspective, however, the strains should be thoroughly tested before industrial use. *T. cantarellii* is also considered a safe yeast for beneficial use according to Bourdichon et al. [Bourdichon et al. 2012].

**Table 4.** Minimum inhibitory concentrations (MIC) of nine antifungal compounds.

| **Antifungal compound** | ***Trigonopsis cantarellii* P-69** | ***Candida sojae* T-39** |
| --- | --- | --- |
| Anidulafungin | 0.12 | 0.25 |
| Amphotericin B | 0.12 | 0.5 |
| Micafungin | 0.12 | 0.03 |
| Caspofungin | 0.12 | 0.06 |
| 5-Flucytosine | >64 | 0.25 |
| Posaconazole | 0.25 | 0.06 |
| Voriconazole | 0.015 | 0.03 |
| Itraconazole | 0.015 | 0.12 |
| Fluconazole | 1 | 1 |

As *T. cantarellii* showed particular promise for low-alcohol beer production, whole-genome sequencing was finally used to confirm the identification of the strain. The paired-end sequencing reads were *de novo* assembled using SPAdes (assembly statistics available in Table 5). As no sequencing data for other *T. cantarellii* strains was found in public databases, the *T. cantarellii* assembly was compared to assemblies of other closely related species [Shen et al. 2018]. These included *Trigonopsis variablis, Trigonopsis vinaria, Sugiyamaella lignohabitans, Tortispora ganteri*, and *Botryozyma nematodophila*. The assembly of *T. cantarellii* P-69 grouped close to *Trigonopsis variabilis, Trigonopsis vinaria* and *Sugiyamaella lignohabitans*, in both a maximum likelihood phylogenetic tree generated based on an alignment of 996 BUSCO genes (Figure 7A), and in a neighbour-joining tree computed based on MinHash (Mash) distances in the assemblies (Figure 7B).

**Table 5.** Assembly statistics for *T. cantarellii* P-69.

|  |  |
| --- | --- |
| Genome size (bp) | 11566117 |
| Contigs | 169 |
| Mean contig size (bp) | 68438 |
| Median contig size (bp) | 2365 |
| N50 | 311069 |
| Largest contig size (bp) | 1002201 |
| GC(%) | 42.88 |
| N count | 696 |
| N(%) | 0.01 |
| Total BUSCO groups searched | 1711 |
| Complete BUSCOs | 960 |
| Complete and single-copy BUSCOs | 956 |
| Complete and duplicated BUSCOs | 4 |
| Fragmented BUSCOs | 188 |

**Figure 7.**
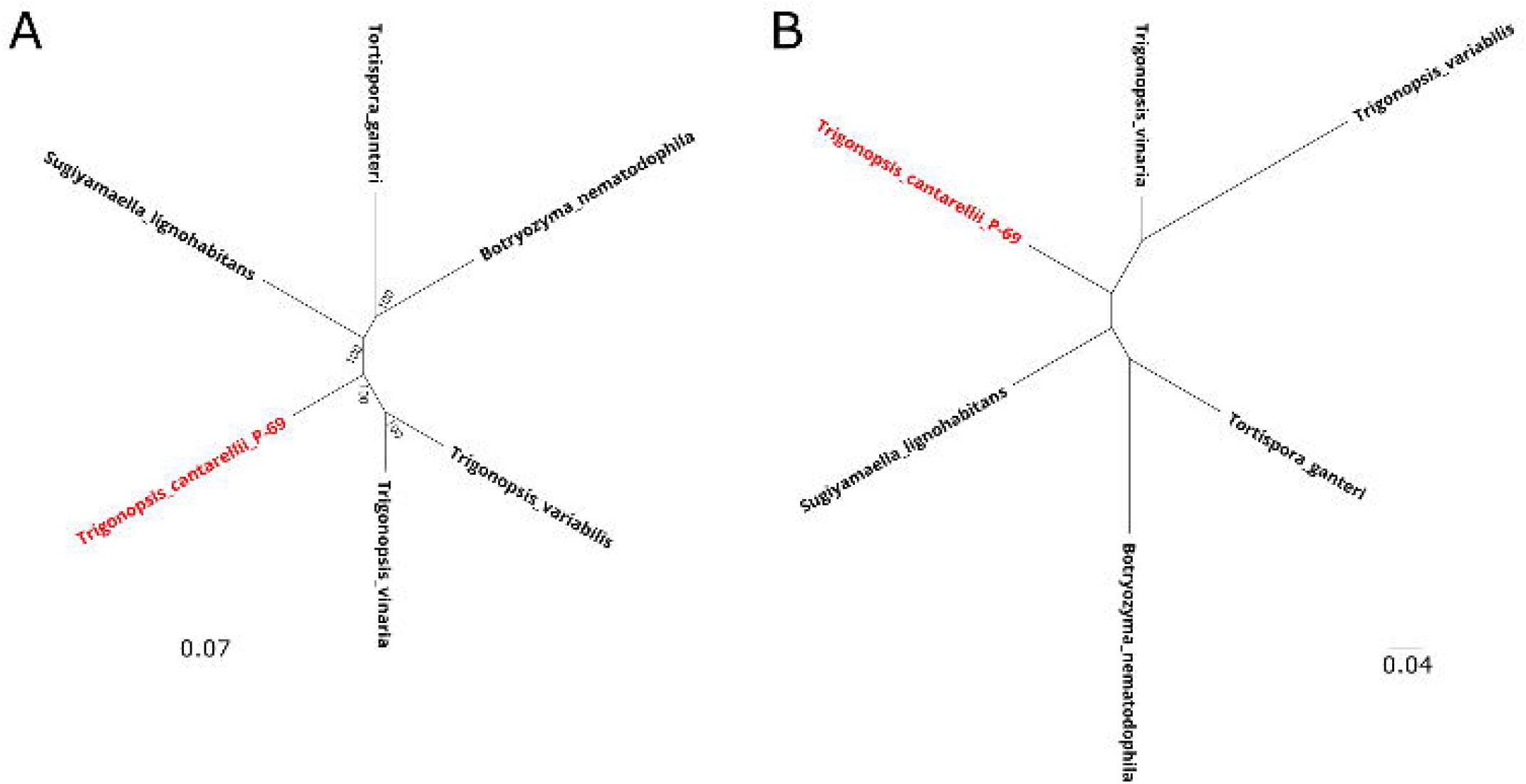
(**A**) A maximum likelihood phylogenetic tree of *Trigonopsis cantarellii* P-69 (coloured red) and five closely related yeast species based on alignment of 996 BUSCO genes. Values at nodes represent bootstrap support. (**B**) A neighbour-joining tree generated based on MinHash distances in the assemblies of *Trigonopsis cantarellii* P-69 (coloured red) and five closely related yeast species.

## Conclusions

In this study we aimed to repurpose and exploit fungal isolates from the natural brewery microbiota for low-alcohol beer production. A number of promising strains for low-alcohol beer fermentation were identified following pre-screening, and two strains were ultimately selected for 30L-scale wort fermentations and descriptive sensory analysis. The two selected strains, *Trigonopsis cantarellii* P-69 and *Candida sojae* T-39, performed comparably to a commercial *Saccharomycodes ludwigii* reference strain. The *T. cantarellii* strain in particular, produced low amounts of off-flavours and a significantly higher amount of the desirable monoterpene alcohol *trans*-geraniol. The mechanisms and potential for monoterpene biotransformation should be studied in more detail in the future. The strain also appears to be easily controllable by common food preservatives in the brewery, and appears to have low risk of pathogenicity.

## Acknowledgements

We thank Aila Siltala, Niklas Fred, Eero Mattila for technical assistance, Atte Mikkelson and Liisa Änäkäinen for the aldehyde analysis, Tuulikki Seppänen-Laakso and Matti Hölttä for the aroma analysis, and Jarkko Nikulin for assistance during sensory analysis. We thank PBL Brewing Laboratory (Oy Panimolaboratorio - Bryggerilaboratorium Ab) and Business Finland for funding the study.

